# Direct Antigen Presentation is the Canonical Pathway of Cytomegalovirus CD8 T-cell Priming Regulated by Balanced Immune Evasion Mounting a Strong Antiviral Response

**DOI:** 10.1101/2023.09.01.555841

**Authors:** Julia K. Büttner, Sara Becker, Annette Fink, Melanie M. Brinkmann, Rafaela Holtappels, Matthias J. Reddehase, Niels A. Lemmermann

## Abstract

CD8 T cells are the main antiviral effectors of the adaptive immune response to cytomegaloviruses (CMVs) in confining acute infection and in long-term surveillance of latent infection. An unresolved issue of debate, with diametrically opposite conclusions, is the mechanism by which CMV-specific naïve CD8 T cells are primed. The idea of priming by antigen cross-presentation through uninfected professional antigen presenting cells taking up viral proteins derived from infected cells was based on the observation that the net response is not improved when direct presentation is enhanced by deletion of viral ‘immune evasion’ genes affecting the classical MHC class-I pathway of antigen presentation. Redundance of priming mechanisms has been demonstrated by experimental models in which either pathway has been rendered inaccessible, so that both pathways are, in principle, capable of priming CD8 T cells. For studying CMV-specific priming in the normal host competent in both antigen presentation pathways, we took the novel approach to enhance instead of delete immune evasion protein expression. Surprisingly, the net magnitude of the CD8 T-cell response in the regional lymph node draining a local site of infection was identical for the contrasts of reduced or enhanced direct antigen presentation. At first glance, this may lead one to conclude that direct presentation plays no role in priming. This interpretation, however, is incompatible with the finding that an intermediate extent of direct antigen presentation, as it exists for wild-type virus, resulted in the best CD8 T-cell response. Our findings thus revealed an essential positive role of a balanced viral immune evasion in mounting a strong antiviral immune response. This novel insight sheds a completely new light on the acquisition of viral immune evasion genes during virus-host co-evolution.

## Introduction

Cytomegaloviruses (CMVs) belong to the β-subfamily of the herpesviruses [reviewed in (1)]. The medical relevance of human CMV (hCMV) is based on its pathogenicity and the resulting multiorgan CMV disease in the absence of immune protection. Risk is associated with congenital infection of the fetus and infection of immunocompromised patients with genetic or acquired immunodeficiencies. Patients with hematopoietic malignancies undergoing hematoablative therapy followed by hematopoietic cell transplantation (HCT) are at risk for developing symptomatic manifestations in the period before full reconstitution. In the case of allogeneic HCT, the risk is further enhanced by immunosuppressive therapy aimed at preventing graft-versus-host disease (GvHD). Likewise, CMV disease poses a threat to recipients of allogeneic solid organ transplantation (SOT) immunosuppressed for preventing graft rejection [reviewed in (2,3)].

The mild and mostly unnoticed infection in the immunocompetent host reflects a largely reduced viral pathogenicity due to the control of viral replication by antiviral effector mechanisms of innate and adaptive immunity. Particularly during transient immunodeficiency in HCT patients, efficient reconstitution of antiviral CD8 T cells is associated with positive prognosis in both clinical CMV infection and in experimental models [(4,5) reviewed in (6)]. Accordingly, adoptive transfer of antiviral CD8 T cells is a promising immunotherapeutic approach to prevent CMV pneumonia and other organ manifestations of CMV infection in HCT recipients (3,7–10).

Due to the strict host-specific replication of CMVs, hCMV cannot be studied in animal models (11,12). As a highly versatile model system of natural host-virus pairs, the infection of mice with murine CMV (mCMV) has been established by many groups for investigating basic principles of viral pathogenesis and antiviral immune control (13–17). These principles are shared between different pairs of CMVs and their respective hosts, as coevolution has led to biological convergences in host-virus adaptation (9).

For both hCMV and mCMV, CD8 T cells have been described as major effectors in preventing viral pathology during acute infection (18,19). In addition, the importance of CD8 T cells in the long-term surveillance of latent mCMV infection (20) is suggested by the expansion of certain viral epitope-specific populations of activated KLRG1^high^ CD62L^low^ CD8 T cells over time, a phenomenon termed ‘memory inflation’ (MI) [for reviews, see (21–24)], and by an enhanced viral transcriptional activity during latency in the absence of such ‘inflating’ CD8 T cells (25). Surprisingly, given the importance of CD8 T cells in the immune control of CMV and the interest in how MI is induced and maintained, the mechanisms underlying CMV-specific priming of naïve CD8 T cells are still not fully understood and remained controversial.

Viral antigenic peptides can be presented in the context of MHC class-I (MHC-I) molecules as pMHC-I complexes by professional antigen-presenting cells (pAPCs) in the lymph nodes to naïve CD8 T cells by two different mechanisms, direct presentation and cross-presentation, both initiating an antiviral immune response. In direct presentation, endogenous viral proteins are processed and presented on infected pAPCs, whereas in cross-presentation uninfected pAPCs take up exogenous antigens and introduce them into the antigen processing and presentation pathway (26). As a potential source of direct antigen presentation for CD8 T-cell priming, mCMV can infect pAPCs *in vitro* and *in vivo*, including dendritic cells (DCs) (27,28) and CD169^+^ macrophages (29).

However, all CMVs code for proteins that manipulate the MHC-I pathway of antigen presentation and are known as viral regulators of antigen presentation (vRAP) [(28), for more recent reviews see (30,31)]. For mCMV, three vRAPs have been described: the positive regulator m04/gp34 and the negative regulators m06/gp48 and m152/gp40. Recent work has shown that m04 and m06, which belong to the same protein family (31), compete for pMHC-I cargo and annihilate each other in their function (32). For this reason, m152 remains as the functionally relevant immunoevasin of mCMV (32) by interfering with the cell surface trafficking of pMHC-I complexes (33–36), thereby reducing their number available for interaction with CD8 T-cell receptors (TCRs) (37). Notably, recent work has shown that a reduction of the number of cell surface pMHC-I molecules raises the avidity threshold required for TCRs of antiviral CD8 T cells to recognize infected cells and protect against infection [reviewed in (38)]. As a consequence, the recognition of infected cells by virus-specific CD8 T cells is impaired by the action of m152 *in vitro* and *in vivo* (28,39,40). Besides downmodulating pMHC-I, m152 has also been shown to interfere with cell surface expression of RAE-1 family ligands of the activating NK cell receptor NKG2D (41–43), thereby preventing NK-cell activation (44,45). Furthermore, m152 target STING to reduce induction of type I antiviral interferons (46). This multifunctionality indicates a key role for m152 in subverting the antiviral defense.

As vRAPs have been shown to be functional in pAPCs after hCMV and mCMV infection (27,28,47–49), thereby interfering with direct antigen presentation, cross-presentation has been hypothesized to be the major pathway of antigen presentation leading to effective CD8 T-cell priming after CMV infections by counteracting viral immune evasion (50–52). Consistent with this, it has been shown that MHC-I-deficient fibroblasts infected with a spread-deficient mCMV, preventing first-round direct presentation and excluding further rounds of infection, can prime mCMV-specific CD8 T cells after immunization of WT mice (53). Furthermore, Batf3^−/−^ mice, which lack cross-presenting CD8α^+^ and CD107^+^ DCs, show impaired priming and MI after mCMV infection (54). All these results clearly indicated that cross-presentation can occur, in principle, during mCMV infection.

On the other hand, there is evidence that priming by cross-presentation does not exclude a role of direct presentation. While Batf3^−/−^ mice completely lack CD8^+^ DCs, which could contribute also to direct priming, infection of CD11c-Rac mice, which are selectively defective in the uptake of exogenous antigens by DCs, showed an mCMV-specific priming comparable to that in WT-mice (51). Similarly, mCMV-specific priming was not impaired in mice treated with the TLR9 agonist CpG to abolish cross-presentation (55). In addition, there is evidence that the downregulation of pMHC-I is less efficient in DCs and macrophages than it is in fibroblasts or endothelial cells (56–58). Overall, it remained controversial whether direct or cross-presentation is the canonical pathway to induce CD8 T-cell responses after CMV infection.

Previous approaches are characterized by blocking either direct antigen-presentation or cross-presentation, and thus were, by concept, unable to decide which pathway is taken with preference in normal mice. Here we introduce the novel approach to study CD8 T-cell priming under conditions of enhanced or prevented immune evasion, leading to reduced or enhanced direct antigen presentation, respectively. In case priming occurs by direct antigen presentation, the outcome has been expected to be diametrically opposed, namely reduced priming after enhanced immune evasion and enhanced priming after prevention of immune evasion. Surprisingly, our data did not reveal such a difference. Rather, the up– or down-modulation of immune evasion gene expression led to the same response magnitude in the net effect. It thus appeared more than logical to conclude that priming is not by direct antigen presentation, thereby providing indirect evidence and an argument for priming by cross-presentation. However, this tempting conclusion did not take into account that modulation of initial antigen presentation has a direct impact on the recognition of infected cells by the newly generated CD8 T cells. In a negative feedback loop, CD8 T cells limit viral replication and thus second-round antigen presentation after initially enhanced antigen presentation. In contrast, poor priming resulting from initially reduced antigen presentation does not limit viral replication and thus leads to enhanced second-round antigen presentation. From this ‘immune evasion paradox’ we conclude that direct antigen presentation is the canonical pathway of mCMV-specific CD8 T-cell priming.

## Materials and Methods

### Cells, viruses, and mice

P815 (No. TIB-64, haplotype H-2^d^) and EL4 (No. TIB-39, haplotype H-2^b^) cells were obtained from the American Tissue Culture Collection (ATCC) and cultivated in RPMI supplemented with 5% fetal calf serum (FCS) and antibiotics, or in DMEM with 10% FCS and antibiotics, respectively. Primary murine embryo fibroblasts (MEF) were cultivated in MEM supplemented with 10% FCS and antibiotics.

A CD8 T-lymphocyte line (CTLL) specific for the immunodominant viral epitope IE1 (YPHFMPTNL) (59) was generated from memory cells derived from latently infected BALB/c mice by four rounds of restimulation of CD8 cells with synthetic peptide (60).

Virus derived from BAC plasmid pSM3fr (61) was used as ‘wild-type’ (WT) virus, mCMV.WT. Recombinant viruses mCMV-Δm152 (34), mCMV-Δm157 (62), and mCMV-m152StopΔm157 (46) have been described previously. BALB/c, C57BL/6, and C57BL/6-Unc93b1^3d/3d^ (3d mice, (63)) mice were bred and housed under specified-pathogen-free conditions at the Central Laboratory Animal Facility (CLAF) at the University Medical Center of the Johannes Gutenberg-University, Mainz, Germany, and at the central animal facility of HZI Braunschweig, Germany. Animal research protocols were approved by the ethics committee of the Landesuntersuchungsamt Rheinland-Pfalz, permission no. 23 177-07/G09-1-004, according to German Federal Law §8 Abs. 1 TierSchG (animal protection law).

### Generation of recombinant viruses

Recombinant plasmids were constructed according to established procedures, and enzyme reactions were performed as recommended by the manufacturers. Throughout, the fidelity of PCR-based cloning steps was verified by sequencing (GATC, Freiburg, Germany).

Mutagenesis of full-length mCMV BAC plasmid pSM3fr (61) was performed in DH10B by using the two-step replacement method as described before (61,64), resulting in the BAC plasmid pSM3fr_m152.IE+E. For this, the shuttle plasmid pST76K_ie2_m152 was used to integrate the open reading frame (ORF) m152 for ectopic expression under the control of the ie2 promoter. In the first step, the intermediate plasmid pST76K_ie1/3-ie2 was generated by subcloning a *Pml*I-cleaved 5,557bp fragment of pUCAMB (65), containing nucleotides 181,415 to 186,972 (GenBank accession no. NC_004065) of the mCMV immediate-early (IE) region into the *Sma*I site of the shuttle plasmid pST76-KSR (64). In a subsequent step, a 1,452bp PCR fragment, covering the ie2 promoter and ORF m152, was introduced in the *Hpa*I cleaved vector to generate pST76K_ie2_m152. The fragment was generated by a touchdown PCR with primer pair m152-*Hpa*I-fwd GAA**GTTAAC**_184,240_CATATAAAAGCTGTCCCCCATGCCATTCGA_184,269-211,468_TCAGACGCGGGCTACTCCCGAAAGAGTAAC_211,439_ and m152-*Hpa*I-rev GGA**GTTAAC**_210,056_TGACTAATAAGTTATCTTTATTGTACAAGTGTTGTGTGTTATCCCTGA GCCCATTCCCAG_210,115_ (*Hpa*I restriction sites are indicated in bold letters) using ProofStart Taq DNA polymerase (catalog no. 202205; QIAGEN, Hilden, Germany) and cycler conditions as follows: an initial step for 5 min at 95°C was followed by 18 cycles for 30 s at 94°C, 120 s with temperatures decreasing by 1°C per cycle starting from 62°C, and 90 s at 68°C each, followed by 12 cycles for 30 s at 94°C, 120 s at 45°C, and 90 s at 68°C.

Reconstitution and purification of a high-titer virus stock of mCMV-m152.IE+E was performed as described (66).

### Infection conditions and virus growth kinetics in immunocompromised mice

For *in vitro* assays, MEF were infected with the indicated viruses at a multiplicity of infection (MOI) of 4 with centrifugal enhancement of infectivity (67,68). Intraplantar infection of 8-to-10-week-old mice was performed in the left hind footpad with 1×10^5^ PFU of the respective virus.

For log-linear *in vivo* virus growth curves, BALB/c mice were immunocompromised by hematoablative treatment with a single 6.5 Gy dose of total-body γ-irradiation and were infected with the respective virus. Quantification of viral genome load in lungs and spleen was performed on days 2, 4, 6, 8, and 10 by qPCR as described previously (66).

### Depletion of lymphocyte subsets *in vivo*

Depletion of NK cells or of CD8 T cells was performed 24 h prior to infection by i.v. injection of 25µl rabbit antiserum directed against asialo-GM1 (catalog no. 986-100001; Wako Chemicals, Osaka, Japan) or of 1mg purified antibody directed against CD8 (clone YTS169.4), respectively (69).

### Quantification of viral genomes and transcripts

To determine viral genome load in lungs and spleen, DNA of infected mice was isolated from the respective tissues with the DNeasy blood & tissue kit (catalog no. 69504; QIAGEN) according to the manufacturer’s instructions. Viral and cellular genomes were quantitated in absolute numbers by M55-specific and pthrp-specific qPCRs normalized to a log_10_-titration of standard plasmid pDrive_gB_PTHrP_Tdy (66,70).

Viral transcripts were quantitated from total RNA, extracted from infected MEF or from lymph nodes, as described previously (69), and 500 ng RNA were used as template for RT-qPCR. Absolute quantification of E1 or m152 transcripts using *in vitro* transcripts as standard has been described previously (71). E1 transcripts were identified for mice of C57BL/6 and BALB/c background, respectively, as a surrogate for viral replication that otherwise would be confounded by inoculum viral DNA. It is important to recall that a PFU of mCMV equals 500 copies of viral genomic DNA (67), so that intraplantar infection with 1×10^5^ PFU corresponds to 5×10^7^ copies, which critically confound the quantitation of *de novo* viral DNA replication in the RLN, in particular at early times. Spliced E1 transcripts (72,73) were chosen as a surrogate, because E1 (M112-113) expression is essential for viral DNA replication (74). It also must be noted that the quantity of E1 transcripts correlates with the number of infected cells.

### Peptides

Custom peptide synthesis to a purity of > 80% was performed by JPT Peptide Technologies (Berlin, Germany). Synthetic peptides representing antigenic peptides in mouse haplotype H-2^b^ were M38 (SSPPMFRVP), M45 (HGIRNASFI), M57 (SCLEFWQRV), M122/IE3 (RALEYKNL), m139 (TVYGFCLL), and m141 (VIDAFSRL) (75). Those for mouse haplotype H-2^d^ were m04 (YPGSLYRRF), m18 (SGPSRGRII), M45 (VGPALGRGL), M83 (YPSKEPFNF), M84 (AYAGLFTPL), M105 (TYWPVVSDI), m123/IE1 (YPHFMPTNL), m145 (CYYASRTKL), and m164 (AGPPRYSRI) (8). The synthetic peptides were used for exogenous loading of target cells in the ELISpot assay.

### ELISpot assay

An interferon gamma (IFNγ) enzyme-linked immunospot (ELISpot) assay was performed for quantification of IFNγ-secreting CD8 T cells after sensitization by peptide-loaded stimulator cells. Frequencies of mCMV-specific CD8 T cells were determined by incubation of graded numbers of immunomagnetically-purified CD8 T cells, derived from the RLN, which is the popliteal lymph node in the case of intraplantar infection, with stimulator cells (P815 or EL4, as it applied) that were exogenously loaded with synthetic peptides at a saturating concentration of 10^−7^M (69). IE1 epitope presentation after endogenous antigen processing in infected BALB/c MEF was determined using short-term IE1-CTLL (76) as responder cells. Spots were counted automatically based on standardized criteria using Immunospot S4 Pro Analyzer (CTL, Shaker Heights, OH, USA) and CTL-Immunospot software V5.1.36. Frequencies of IFNγ-secreting cells and the corresponding 95% confidence intervals were calculated by intercept-free linear regression using Mathematica, version 8.0.4.

### IE phase arrest of infected cells

For a selective arrest of viral gene expression in the IE phase, MEFs were infected in the presence of 50µg/ml cycloheximide (CHX) to block protein synthesis reversibly. At 3 h after infection, the culture medium was replaced by fresh medium containing 5 µg/µl actinomycin D (ActD) as described previously (77).

### Intracellular cytokine assay

1×10^5^ mCMV-infected, IE-arrested MEFs (BALB/c, haplotype H-2^d^) were co-cultivated with 5×10^5^ IE1-CTLL cells for 5h at 37°C in the presence of brefeldin A (BD GolgiPlug, final concentration 1:1,000; catalog no. 555029; BD Biosciences). Subsequently, the CTLL cells were fixed, permeabilized with BD Cytofix/Cytoperm (catalog number 554722, BD Biosciences) and stained with FITC-conjugated anti-mouse IFNy antibody (clone XMG1.2, catalog no. 554411, BD Biosciences).

### Genome-wide ORF library screening

An mCMV ORF library of expression plasmids spanning the whole mCMV genome (75) was used for ORF-specific stimulation of *ex vivo* isolated CD8 T cells with transfected SV-40 fibroblasts followed by cytofluorometric detection of intracellular IFNγ.

### Immunoblot analysis

The expression of mCMV proteins was detected by Western blot analysis as described (34). In brief, lysates of infected MEF were prepared and 30µg of total protein amount was subjected to separation by 12.5% SDS-PAGE, followed by blotting onto polyvinylidene difluoride membrane and protein labeling with the respective antibodies. The following antibodies were used: m152 (M3D10, monoclonal antibody, kindly provided by E. Kremmer, Helmholtz Zentrum München, Munich, Germany), IE1 (Croma 101, monoclonal antibody, kindly provided by S. Jonjic, University of Rijeka, Rijeka, Croatia), and IE2 (αIE2-N, polyclonal antibody, rabbit, Peptide Specialty Laboratories, Heidelberg, Germany). Detection of antibody binding was visualized by chemiluminescence using the ECLplus Western blotting detection system (catalog no. RPN2132; Amersham Bioscience, Little Chalfont, United Kingdom) and Lumi-Film (catalog no. 11666657001, Roche Applied Science, Mannheim, Germany).

### Statistical analysis

To evaluate statistical significance of differences between two independent sets of data, the unpaired t-test with Welch’s correction of unequal variances was used. Differences were considered statistically significant for p values <0.05 and highly significant for <0.001.

Virus doubling times (vDT) and their 95% confidence intervals (CI) were calculated from the slopes of log-linear regression lines determined by linear regression analysis (78–80). Calculations were performed with Graph Pad Prism 6.04 (Graph Pad Software, San Diego, CA). It should be noted that vDT values vary between different organs but are a constant for each organ independent of the viral replication parameter tested, that is, identical for viral genomic DNA copy numbers measured by qPCR, infectious virus expressed as PFU, or numbers of infected tissue cells determined by immunohistological detection of viral proteins (79).

## Results

### Broad CD8 T-cell response to mCMV in C57BL/6 mice genetically deficient in the antigen cross-presentation pathway

Conclusions on the mechanism of CD8 T-cell priming are usually based on measuring the magnitude of the primary immune response. This follows the tacit assumption that antigen presentation requirements are identical for sensitization of naïve CD8 T-cells by antigen presentation at the immunological synapse (81,82), which is the initial priming event, and subsequent clonal expansion. Accordingly, the terms ‘priming’ and ‘primary response’ are usually used as synonyms, not least because the initial priming event is difficult to access. Thus, while a response definitely indicates a successful initial priming, one must keep in mind that one actually looks at the combined result of priming and subsequent clonal expansion.

To provide evidence for or against cross-priming, we studied here mCMV-specific priming in a regional lymph node (RLN) draining a site of local infection, which is the anatomical structure where direct priming is said to occur (83). Our work builds on a previous own study in which we have shown that mCMV rapidly reaches the RLN that drains the site of infection, infects cells in the peri-subcapsular sinus region, and generates virus-specific effector CD8 T cells as early as by day 3 after virus exposure (69). Notably, this finding is in perfect accordance with the more recent study by Reynoso and colleagues (84) showing that lymph node conduits rapidly transport virions to infect cells in the RLN paracortex for rapid direct T-cell priming in the T-cell zone.

Accordingly, with the assumption that direct priming applies as proposed (83), we analyzed the mCMV-specific CD8 T-cell response in Unc93b1^3d/3d^ mice (Fig. 1). This strain of C57BL/6 genetic background is known to lack endosomal TLR3, 7, and 9 signaling and is impaired in exogenous antigen processing, resulting in increased susceptibility to infection and blockade of cross-presentation (63,85–88). Previous reports on mCMV infection in these mice revealed impaired cytokine production but comparable frequencies of M45–specific hepatic CD8 T cells (63,89). Specifically, we compared the CD8 T-cell response in C57BL/6 WT and Unc93b1^3d/3d^ mice not just for M45 but for a panel of known mCMV peptides presented in the H2^b^ haplotype (Fig. 1). Overall, no qualitative differences in the immunodominance patterns were found. To our surprise, the magnitude of the CD8 T-cell response to many of the tested peptides, most obvious for M45, m139, and m141, was even higher in cross-presentation-deficient Unc93b1^3d/3d^ mice, an observation consistent with previous findings for mice in which cross-presentation was suppressed by CpG pretreatment (55).

**Fig. 1:**
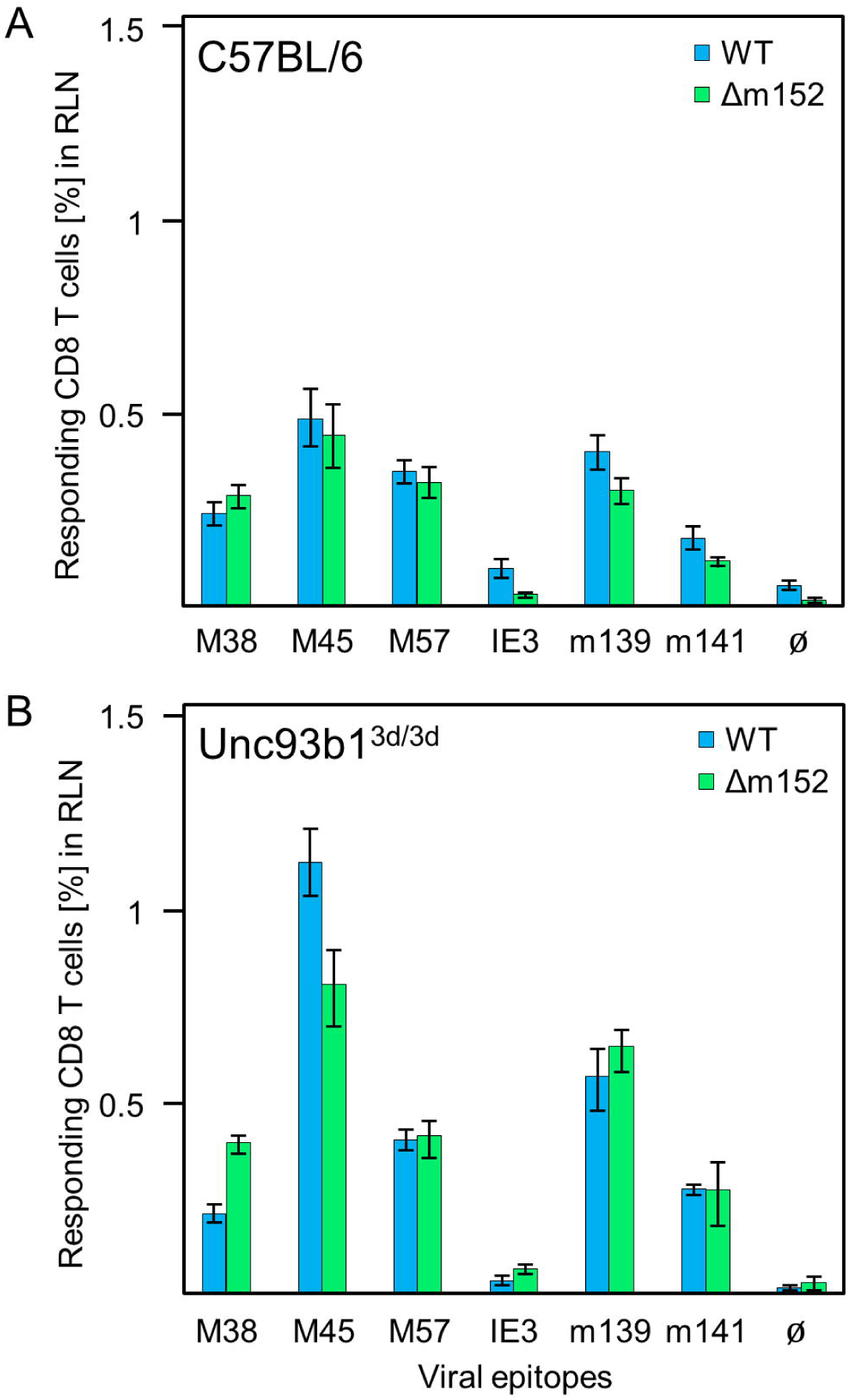
CD8 T-cell response induced by mCMV in presence and absence of the cross-priming pathway. (A) CD8 T-cell response in C57BL/6 mice. (B) CD8 T-cell response in C57BL/6-Unc93b1^3d/3d^ mice that are deficient in antigen cross-presentation. CD8 T cells were isolated from the draining regional lymph node (RLN), that is, the popliteal lymph node, on day 7 after intraplantar infection with 1×10^5^ PFU each of either mCMV.WT (WT) or mCMV-Δm152 (Δm152), and used as effector cells in an IFNγ-based ELISpot assay. EL4 cells exogenously loaded with the indicated synthetic peptides at a saturating concentration of 10^−7^ M were used as APCs. Bars represent most probable numbers (MPN) of responding cells determined by intercept-free linear regression analysis, and error bars represent the corresponding 95% confidence intervals. Ø, no peptide added.

In addition, we analyzed the response under conditions of enhanced direct antigen presentation by infecting both mouse strains with mCMV-Δm152. Here, the infection resulted in essentially the same response magnitude as seen after mCMV.WT infection. These results definitely demonstrated that mCMV-specific priming is possible in the absence of cross-presentation, though, surprisingly, also regardless of whether or not direct antigen presentation was inhibited by the most effective mCMV vRAP m152. All in all, the CD8 T-cell response to mCMV infection turned out to be remarkably resilient and buffered.

### An early NK-cell response simultaneously restricts intranodal viral replication and the CD8 T-cell response in C57BL/6 mice

We have reported previously for the immune response in BALB/c mice that deletion of vRAPs influences the response magnitude in a negative feedback loop by limiting intranodal viral replication. This reduces the supply of antigen needed for the expansion of recently primed CD8 T cells, thereby resulting in a lower response magnitude. As one would have expected just the opposite result, namely an enhanced response resulting from improved direct antigen presentation in the absence of vRAPs, we called this phenomenon the ‘immune evasion paradox’ (69).

In contrast to these own previous findings in BALB/c mice, the mCMV-specific response in C57BL/6 mice is not notably affected by vRAP expression (Fig. 1A). This has originally been shown for the M45 epitope by Gold et al. (90). One difference between the two mouse strains in an early control of viral replication due to the strong activation of Ly49H^+^ NK cells by the viral protein m157 selectively in C57BL/6 mice (91,92), as BALB/c mice do not express Ly49H and accordingly lack this subset of NK cells. It has also been shown that the activation of Ly49H^+^ NK cells by m157 suppresses the mCMV-specific CD8 T-cell response (93). Thus, Ly49H^+^ NK cells likely are a player that contributes to the ‘buffered response magnitude’ in C57BL/6 mice.

For clarification, we investigated the impact of NK cells on the CD8 T-cell response in the draining RLN measured on day 7 after intraplantar infection of C57BL/6 mice, and correlated the response magnitude with viral replication at this site on day 3 (Fig. 2). In the presence of NK cells and CD8 T cells, mCMV.WT and mCMV-Δm152 induced similar CD8 T-cell responses by day 7 (Fig. 2A1), consistent with the preceding experiment (Fig. 1A). Notably, transcription of viral gene E1, which serves as a surrogate for viral replication, was identical for both viruses and on a low level (Fig. A2), so that recently primed CD8 T cells apparently could not further reduce the anyway low viral replication by negative feedback. In contrast, depletion of pan-NK cells prior to infection led to an increase in the response magnitude, at least for the more immunodominant viral peptides (M45, M57, and m139), but only when direct antigen presentation was enhanced by deletion of m152 (Fig.2B1). As it was predicted, depletion of NK cells led to a greatly enhanced viral replication in the RLN (Fig. 2B2) compared to the control group with no NK-cell depletion (Fig. 2A2). Deletion of m152 resulted in a slight, but statistically significant, reduction in viral replication compared to WT virus infection (Fig. 2B2). This reduction was obviously caused by recently primed CD8 T cells, as the difference was abolished by the depletion of CD8 T cells (Fig. 2C).

**Fig. 2:**
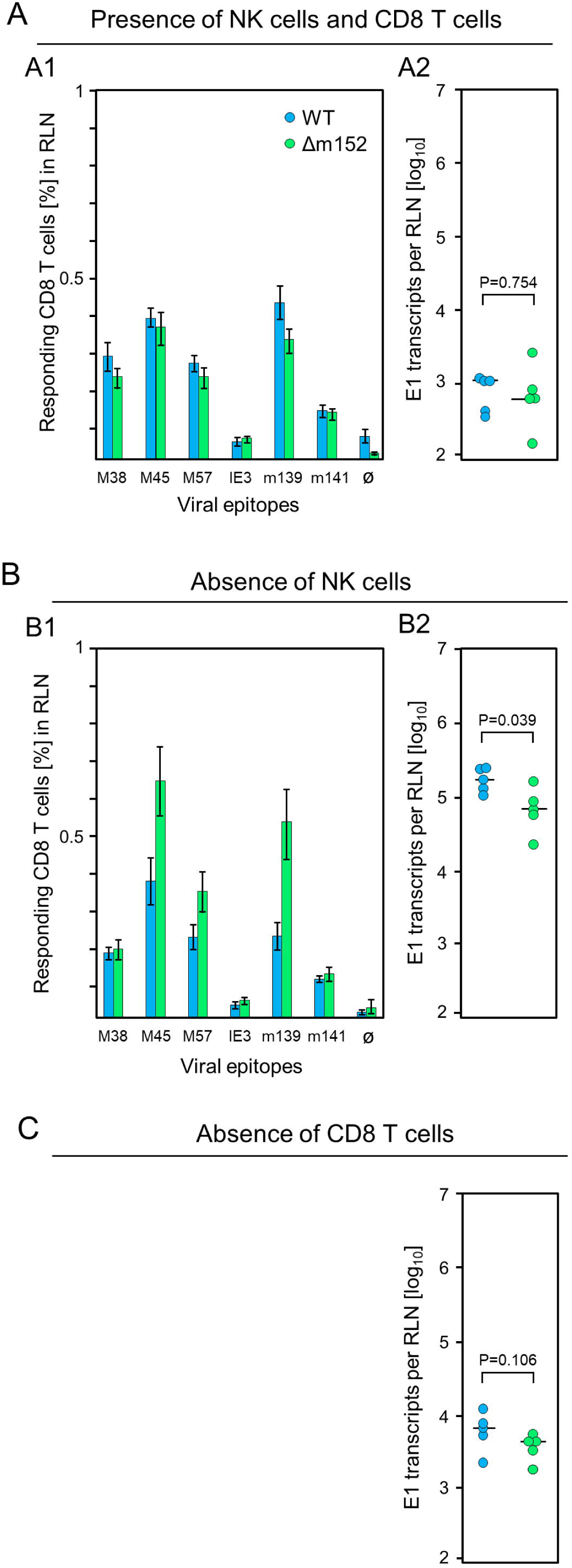
Influence of lymphocyte subsets on the CD8 T-cell response and the intranodal viral replication in C57BL/6 mice. Intraplantar infection of immunocompetent C57BL/6 mice was performed with 1×10^5^ PFU each of either mCMV.WT (WT) or mCMV-Δm152 (Δm152). (A) No depletion of lymphocyte subsets. (B) Depletion of NK cells one day before infection. (C) Depletion of CD8 T cells one day before infection. The CD8 T-cell response in the RLN on day 7 post-infection was assessed by an IFNγ-based ELISpot assay (A1, B1), as explained in greater detail in the legend of Fig. 1. As a surrogate for viral replication in the presence of otherwise confounding viral inoculum DNA, spliced E1 transcripts present in the RLN were quantitated by RT-qPCR on day 3 post-infection (A2, B2, C). Symbols represent individual mice and horizontal bars indicate median values. P values were calculated on the basis of the log-transformed data with Welch’s unpaired t-test (two-sided).

These findings were essentially reproduced in an independent additional experiment using the alternative approach of testing the influence of m152 on the response magnitude in infected C57BL/6 mice in the absence specifically of Ly49H^+^ NK-cell activation via Ly49H-m157 ligation, instead of pan-NK cell depletion (Fig. 3). For this, we compared response magnitude and viral replication in the RLN after infection with the viruses mCMV-Δm157, expressing m152, and mCMV-m152StopΔm157, lacking m152 expression, both in the absence of Ly49H^+^ NK cell activation (Fig. 3). Notably, deletion of m152 increased the CD8 T-cell frequencies for the antigenic peptides M45, M57, and m139 (Fig. 3A1), almost paralleling the findings after pan-NK cell depletion. Under these conditions, too, early recognition of infected cells is indicated by a reduction in viral replication in the RLN after infection with mCMV-m152StopΔm157 (Fig. 3A2). Again, this antiviral control in the absence of Ly49H^+^ NK-cell activation is mediated by recently primed CD8 T cells, as depletion of CD8 T cells led to an increase in viral replication in the RLN (Fig. 3B compared to Fig. 3A2).

**Fig. 3:**
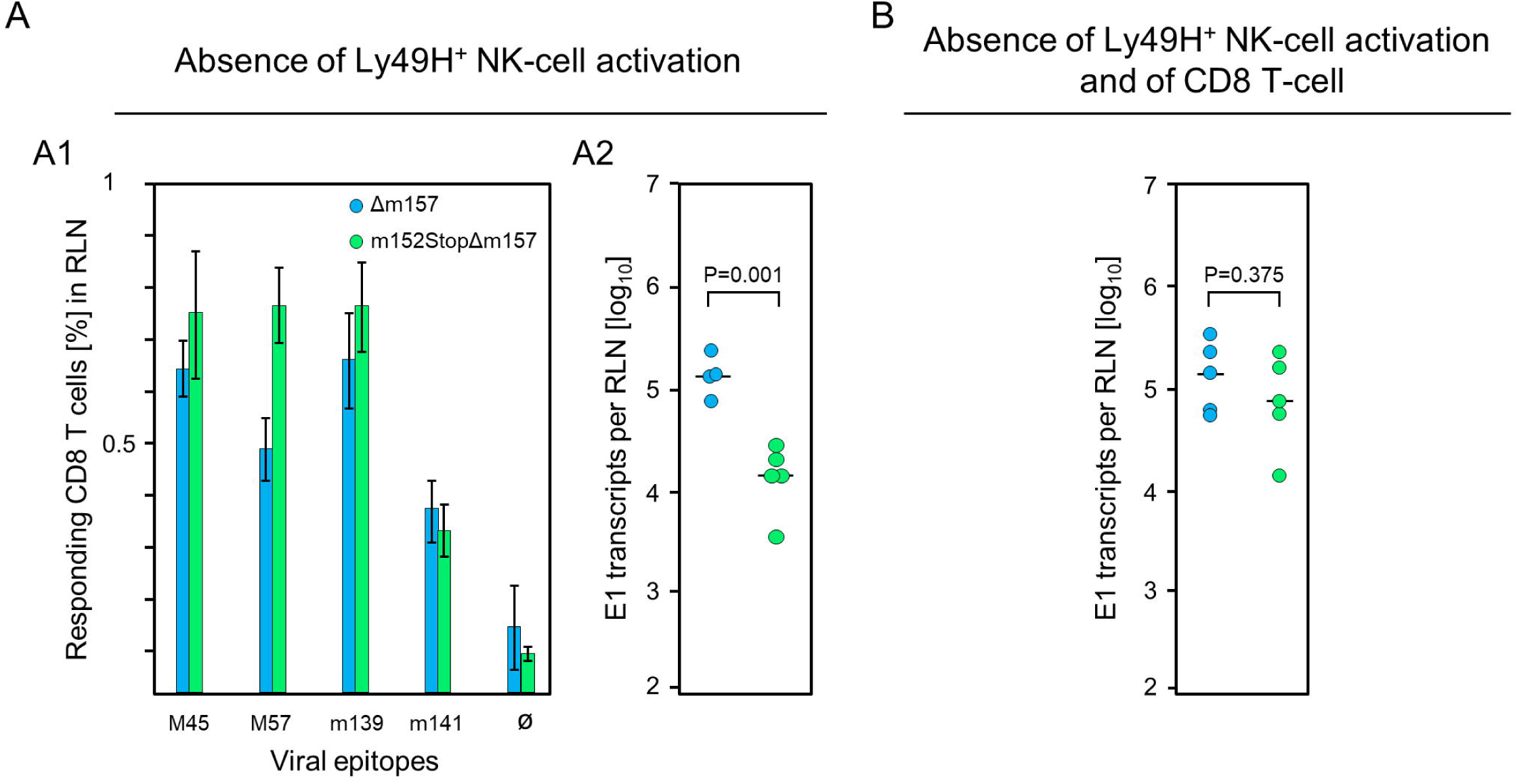
Impact of Ly49H^+^ NK cells on CD8 T-cell response and intranodal viral replication in C57BL/6 mice. (A) Absence of Ly49H^+^ NK-cell activation. (B) Absence of Ly49H^+^ NK-cell activation and of CD8 T cells. Intraplantar infection of immunocompetent C57BL/6 mice was performed with 1×10^5^ PFU each of either mCMV-Δm157 (Δm157) or mCMV-m152StopΔm157 (m152StopΔm157). (A1) The CD8 T-cell response in the RLN on day 7 post-infection was assessed by an IFNγ-based ELISpot assay, as explained in greater detail in the legend of Fig.1. (A2, B) Spliced E1 transcripts present in the RLN on day 3 post-infection were quantitated by RT-qPCR. Depletion of CD8 T cells in (B) was performed on the day before infection. Symbols represent data from individual mice and horizontal bars indicate median values. P values were calculated on the basis of the log-transformed data with Welch’s unpaired t-test (two sided).

Taken together, our results are consistent in having demonstrated that early viral replication in the RLN of C57BL/6 mice is mainly controlled by Ly49H^+^ NK cells, which are activated through Ly49H-m157 ligation. However, unlike the negative feedback loop described for BALB/c mice to be mediated by recently primed CD8 T cells that limit antigen supply for a potential cross-presentation, resulting in a reduced net response (69), reduction of viral replication in the RLN of C57BL/6 mice did not correspond to a lower but to a higher CD8 T-cell response in absence of m152. These data thus clearly support the hypothesis of direct antigen presentation.

All in all, prevention of immune evasion did not lead to a clear picture of whether direct or cross-presentation dominates mCMV-specific priming regardless of genetic differences between mouse strains. Specifically, while all data shown here argued for direct antigen presentation prevailing in mice of C57BL/6 background, cross-presentation was previously thought to contribute to the CD8 T-cell response in BALB/c mice. Thus, a new approach was needed to clarify the mode of antigen presentation in BALB/c mice.

### Combined kinetic acceleration and enhancement of immune evasion strongly inhibit direct antigen presentation

As a new approach for studying the mechanism of mCMV-specific CD8 T-cell priming in BALB/c mice, we enhanced immune evasion both by earlier onset and increased expression of the major immune evasion protein m152. In mCMV.WT infection, m152 is expressed starting very early in the Early (E) phase of the viral replication cycle (33,94). Thus, antigens expressed earlier, that is, in the Immediate-Early (IE) phase, may profit from a head start advantage of presentation before immune evasion operates.

As a new study tool, we constructed the recombinant virus mCMV-m152.IE+E that expresses m152 in both the IE phase and from the E phase onward. Ectopic expression in the IE phase was achieved by insertion mutagenesis placing ORF m152 under the control of the IE2 enhancer-promoter, thereby disrupting the IE2 gene. Expression as an E phase protein occurred from its authentic position in the mCMV genome (Fig. 4A). For verifying immune evasion operating already in the IE phase, viral gene expression was metabolically arrested in the IE phase (Fig. 4B1). This led to an enhanced synthesis of IE proteins IE1 and IE2, and absence of the m152 protein, after infection with either mCMV.WT or the deletion mutant mCMV-Δm152, whereas all three glycosylation isoforms of m152 (34) were strongly expressed and IE2 was absent after infection with mCMV-m152.IE+E (Fig. 4B2). Presentation of the antigenic peptide IE1 (YPHFMPTNL, presented by L^d^) on infected BALB/c-derived cells was tested functionally with an IE1 epitope-specific CTLL (IE1-CTLL) used as responder cells in an IFNγ-based ELISpot assay with IE phase-arrested infected cells as stimulator cells (Fig. 4B3) as well as by intracellular cytofluorometric staining of IFNγ in IE1-CTLL cells sensitized by IE phase-arrested infected cells (Fig. 4B4). In both read-out assays, IE1-peptide was presented after infection with either mCMV.WT virus or the deletion mutant mCMV-Δm152, whereas presentation was blocked after infection with mCMV-m152.IE+E (Fig. 4B3, and 4B4).

**Fig. 4:**
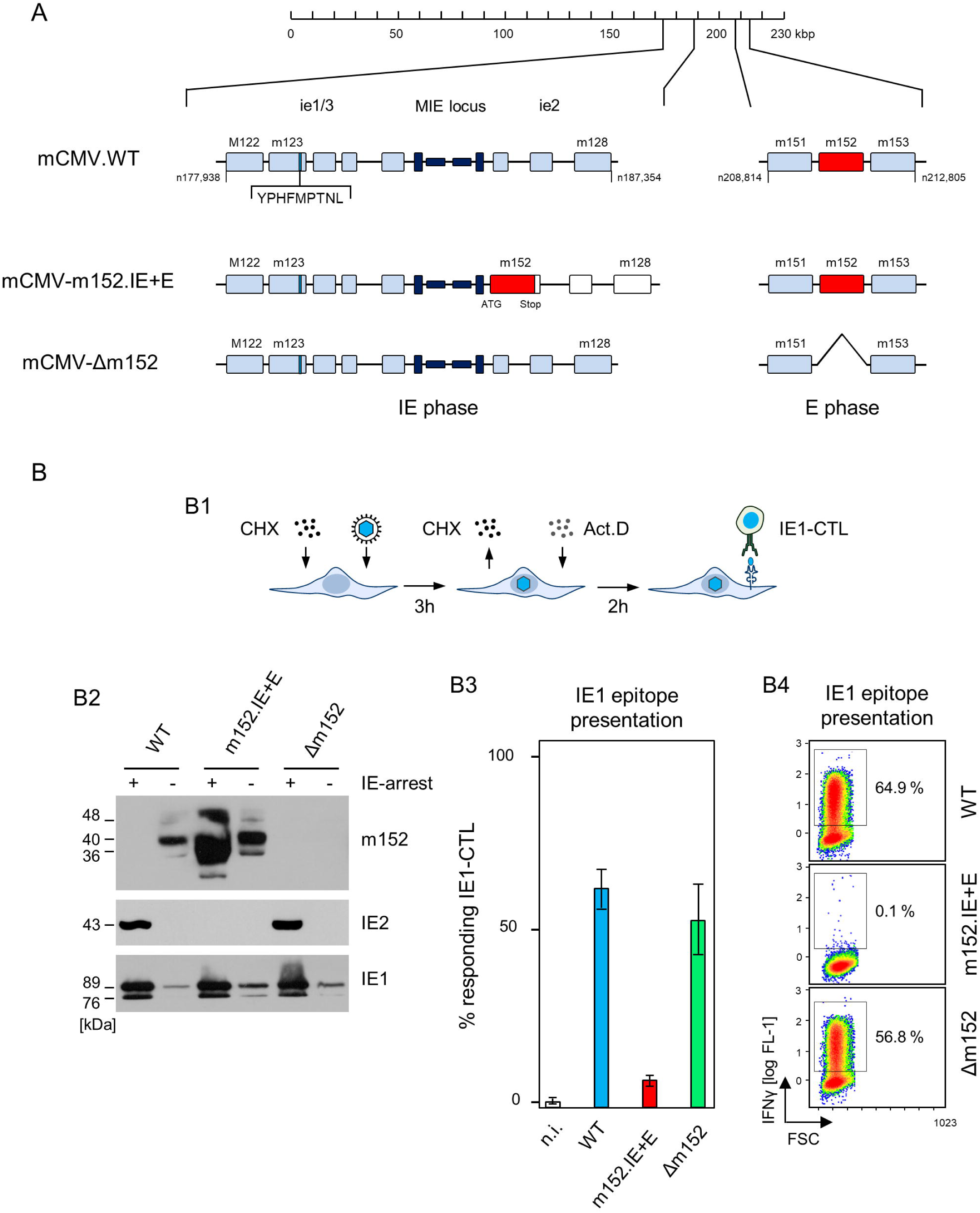
Construction and *in vitro* characterization of recombinant virus mCMV-m152.IE+E. (A) Maps, illustrating the mutagenesis design. A map of the mCMV genome is shown at the top of the sketch. The genomic region harboring the parental E phase gene m152 is shown expanded on the right-hand side. The Major Immediately Early (MIE) locus, consisting of the MIE promotor-enhancer-enhancer-promotor (dark blue) flanked by transcription units ie1/3 and ie2, is shown expanded on the left-hand side. Exons are indicated by boxes (light blue). Recombinant virus mCMV-m152.IE+E, expressing m152 in both the IE phase and the E phase, was generated by introducing the ORF m152 by two-step BAC mutagenesis into the MIE locus under the control of the IE2 enhancer-promoter element, thereby disrupting the IE2 gene. (B) Immune evasion under selective IE phase conditions. (B1) Experimental protocol for arresting infection in the IE phase. BALB/c-derived mouse embryonal fibroblasts (MEF) were infected in the presence of cycloheximide (CHX) that was replaced at 3 h post-infection with actinomycin D (ActD). The thus IE phase-arrested MEF were harvested for analyses at 5 h post-infection. (B2) Western blot analysis of m152 (40kDa and additional glycosylaton isoforms), IE2 (43kDa) and IE1 (89/76kDa) protein expression in IE phase-arrested (+) or untreated (-) MEF infected with the indicated viruses. (B3) IFNγ-based ELISpot analysis quantifying cells of an IE1-CTLL that were sensitized by IE phase-arrested BALB/c-derived MEF infected with the indicated viruses. Bars represent most probable numbers (MPN) of responding cells determined by intercept-free linear regression analysis, and error bars represent the corresponding 95% confidence intervals (n.i) not infected. (B4) Intracellular IFNγ-staining of IE1-CTLL cells at 5 h after co-cultivation with IE phase-arrested MEF infected with the indicated viruses. Two-dimensional color-coded density plots (with red and blue representing highest and lowest density, respectively) show intracellular IFNγ expression (ordinate; fluorescence intensity) versus the forward scatter (abscissa; FSC, linear scale of channels) with 50,000 cells analyzed. The percentages of IFNγ^+^ cells present in the demarcated gates are indicated.

As infected APCs that prime naïve CD8 T cells *in vivo* are not usually arrested in the IE phase, we studied the kinetics of m152 transcription in cells infected with either mCMV.WT, expressing m152 only in the E phase and onward, or mCMV-m152.IE+E, expressing m152 additionally already in the IE phase (Fig. 5A). The transcription data were then correlated with presentation of the antigenic IE1 peptide detected by sensitization of IE1-specific CTLL cells (Fig. 5B). In accordance with the underlying concept of constructing the mutant virus mCMV-m152.IE+E, m152 was expressed faster and inhibited antigen presentation earlier compared to infection with mCMV.WT. In fact, at the early and late time of the infection kinetics, the IE1 peptide was not detectably presented by cells infected with mCMV-m152.IE+E. In contrast, under conditions of ‘physiological’ immune evasion gene expression, as it applies to infection with mCMV.WT, the IE1 peptide was presented early in the time course but absent later (Fig. 5B). We thus conclude that direct antigen presentation is largely inhibited after infection with mCMV-m152.IE+E.

**Fig. 5:**
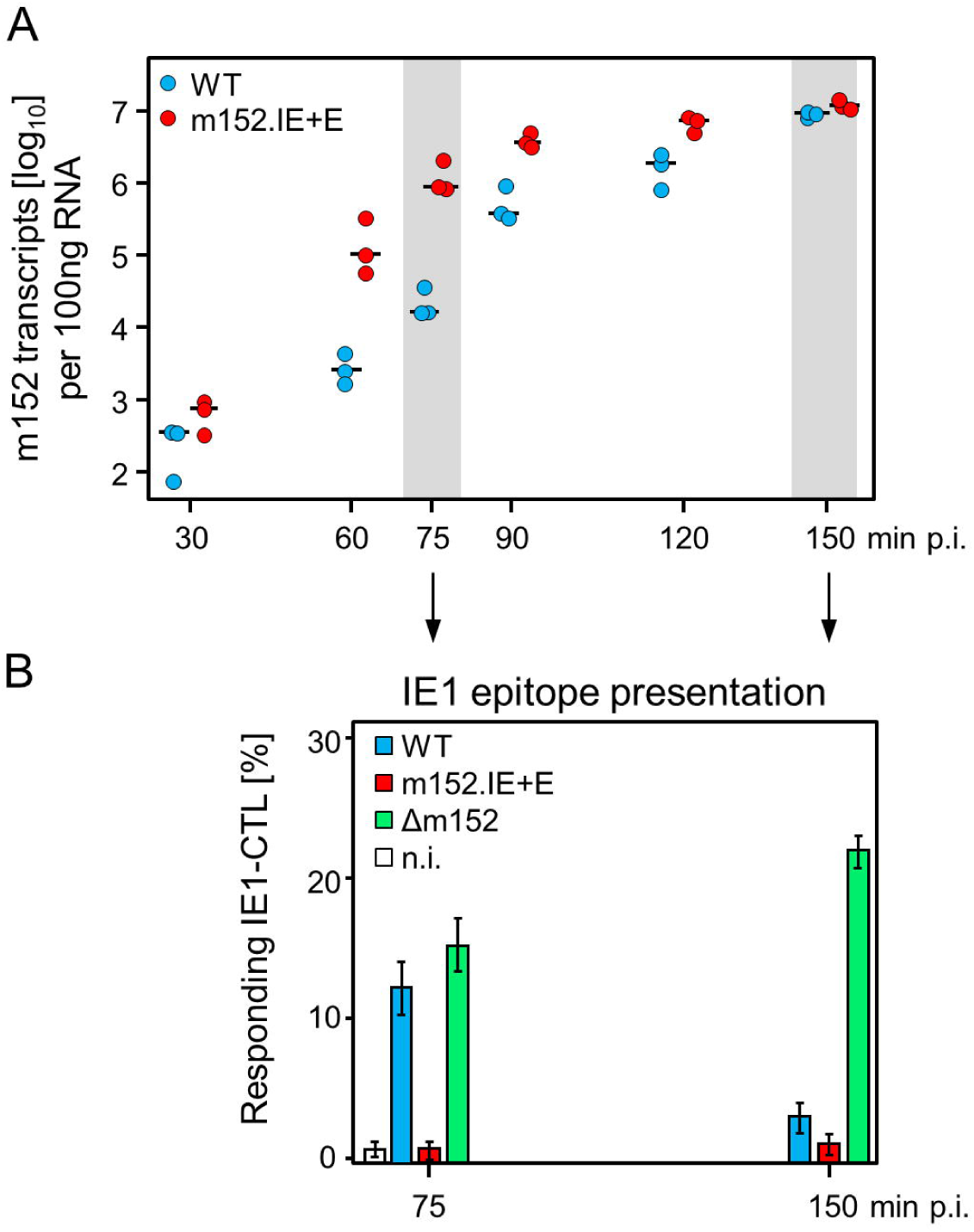
mCMV-m152.IE+E infection inhibits IE1 peptide presentation from the start of viral gene expression. (A) Quantitation of m152 transcripts in the time-course. BALB/c-derived MEF were infected with the indicated viruses and m152 transcripts were quantitated by RT-qPCR, normalized to 100 ng of total RNA, at the indicated times post-infection. Symbols represent biological replicates. Median values are indicated by horizontal bars. (B) IE1-peptide presentation at selected times. IE1-peptide presentation was detected at the indicated times based on sensitization of IE1-CTLL cells in an IFNγ-based ELISpot assay by BALB/c-derived MEF infected with the viruses indicated. To avoid ongoing transcription in the stimulator cells during the ELISpot assay time, the transcription inhibitor ActD was added at the indicated times of cell harvest. Bars represent the percentage of responding cells, and error bars indicate the 95% confidence intervals determined by intercept-free linear regression analysis. (n.i.) no infection.

### Enhanced immune evasion restricts the CD8 T-cell response in BALB/c mice despite an increased viral replication delivering a higher amount of antigen available for cross-presentation

BALB/c mice are competent in both antigen presentation pathways and therefore have the choice. This raised the question of which pathway is used as the canonical pathway for priming of naïve CD8 T cells and for subsequent clonal expansion or whether the pathways replace each other in case of need.

Assuming that direct antigen presentation is the preferred mode of priming, one expects a response magnitude in a rank order defined by the quantity of pMHC-I complexes on the surface of infected APCs and thus reciprocal to the strength of immune evasion: that is, highest response after infection with mCMV-Δm152, intermediate response after infection with mCMV.WT, which corresponds to mCMV-m152.E, and lowest response after infection with mCMV-m152.IE+E. The results of the experiment posed us a riddle. Consistently for a panel of antigenic peptides, and thus reproducibly, the response magnitude was essentially the same for the extremes of highest and lowest direct antigen presentation after mCMV-Δm152 and mCMV-m152.IE+E infection, respectively (Fig. 6A). In linear thinking, the immediate and easiest explanation is that irrelevance of the degree of immune evasion excludes a role for direct antigen presentation. However, the results for mCMV.WT do not fit the mold, as the CD8 T-cell response to this virus was higher in terms of magnitude (Fig. 6A, Suppl. Fig. 1) and broader in terms of the epitope specificity repertoire (Suppl. Fig. 1) when compared to the two mutant viruses. So, if direct presentation would play no role at all, as it was suggested by the two antipodal mutants, why is the response best after infection with mCMV.WT, for which immune evasion and direct antigen presentation are intermediate?

**Fig. 6:**
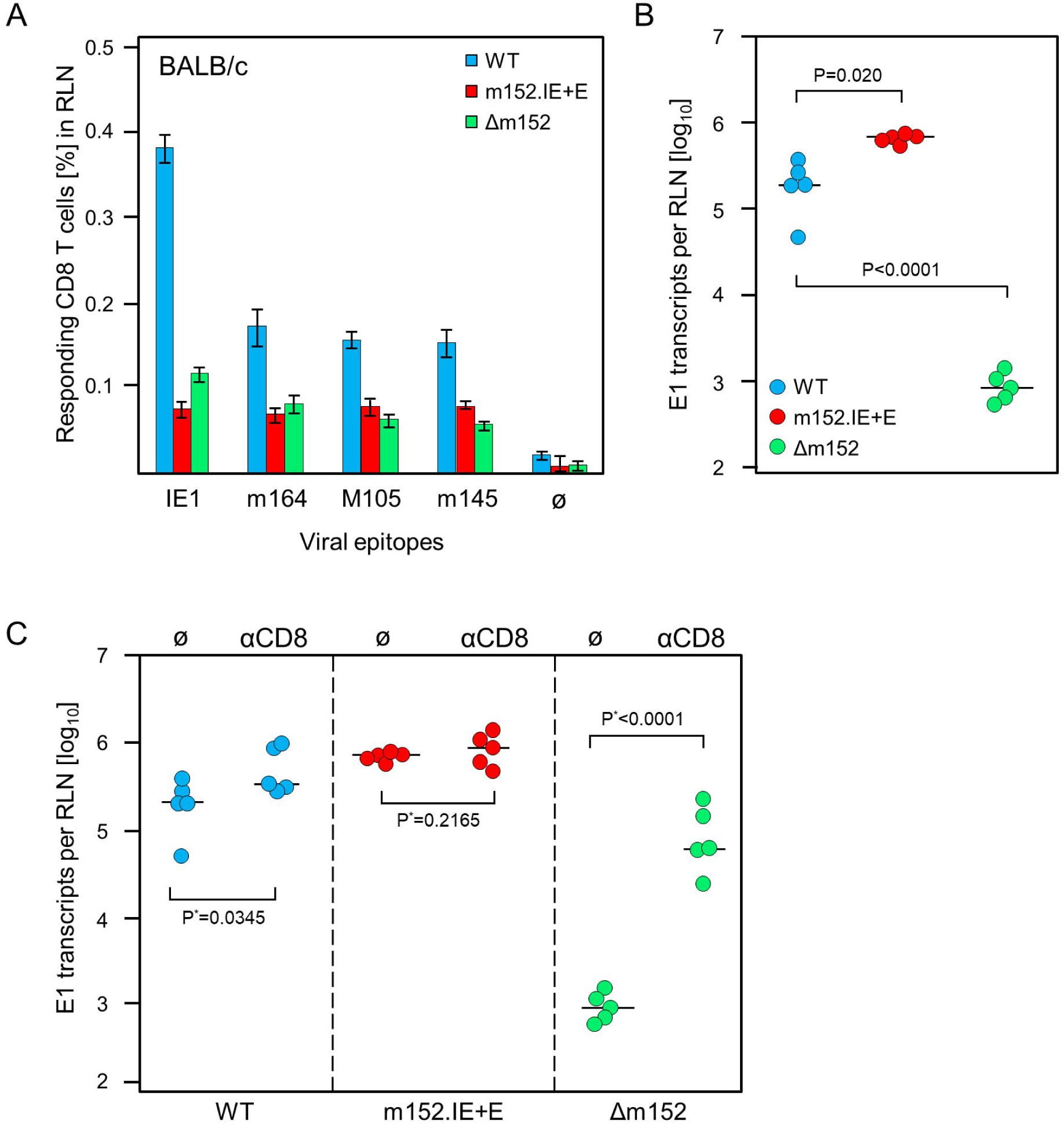
Impact of the strength of viral immune evasion on CD8 T-cell response magnitude and control of intranodal viral replication. Intraplantar infection of BALB/cmice was performed with 1×10^5^ PFU each of either mCMV.WT (WT) or mCMV-m152.IE+E (m152.IE+E) or mCMV-Δm152 (Δm152). (A) CD8 T-cell response in the draining RLN on day 7 post-infection. Responding CD8 T cells were quantitated in an IFNγ-based ELISpot assay, with P815 cells exogenously-loaded with the indicated synthetic peptides as APCs. Bars represent frequencies of epitope-specific CD8 T cells determined by intercept-free linear regression analysis, error bars indicate the 95% confidence intervals. Ø, no peptide added. (B) Viral replication in the RLN on day 3 post-infection, determined by quantitating spliced E1 transcripts by RT-qPCR. Dots represent data for individual mice. Median values are indicated by horizontal bars. (C) Intranodal viral replication in mice depleted of CD8 T cells on the day before intraplantar infection (αCD8) or in mice left undepleted under otherwise identical conditions (Ø). Note that data for undepleted mice are the same as in panel B and are shown again to facilitate the comparison with results from mice depleted of CD8 T cells. P-values are calculated from log-transformed data using Welch’s unpaired t-test (one-sided).

Based on our experience in the C57BL/6 model (Fig. 2 and Fig. 3) and our previous work in the BALB/c model (69) we quantitated replication of the three viruses in the draining RLN, the site where priming and clonal expansion take place, on day 3 after infection (Fig. 6B). As an important control, replication differences reflecting differences in the genetically-determined viral replicative fitness were excluded by showing identical replication of the three viruses in immunodeficient mice (Suppl. Fig. 2). Hence, replication differences in the RLN reflect immune control.

Notably, unlike the response magnitudes, the intranodal viral replication fulfilled the logic, that is, highest replication corresponds to strongest immune evasion for mCMV-m152.IE+E, lowest replication corresponds to weakest immune evasion for mCMV-Δm152, and intermediate replication corresponds to intermediate immune evasion for mCMV.WT (Fig. 6B).

The contribution of recently primed virus-specific CD8 T cells to the control of intranodal virus replication was tested by CD8 T-cell depletion prior to infection (Fig. 6C). It is important to note that our previous work has already shown that antiviral effector CD8 T cells are present by day 3 after infection (69). In line with almost missing antigen presentation on cells infected with mCMV-m152.IE+E, intranodal viral replication was not detectably controlled by CD8 T cells, whereas, in accordance with optimal antigen presentation on cells infected with mCMV-Δm152, intranodal viral replication was almost prevented when CD8 T cells were present. Again, results for mCMV.WT were in between.

In conclusion, diametrically different levels of intranodal viral replication of mCMV-m152.IE+E and mCMV-Δm152 reflect missing and particularly strong antiviral control, respectively, by recently primed CD8 T cells. Astonishingly, the net result is almost the same low response magnitude in the end, whereas the intermediate strength of immune evasion by mCMV.WT results in the best response, generating high numbers of antiviral CD8 T cells for export from the RLN to control virus replication at distant organ sites of viral pathogenesis.

## Discussion

CMVs are often discussed as being masters in evading the innate and adaptive immune control (30,52,95), whereas host counter-measures to ensure immune surveillance of CMV were rarely considered (96). Numerous reports published over decades specifically dealt with viral proteins, so called immunoevasins, which subvert the MHC-I pathway of direct antigen presentation to CD8 T cells [for reviews, see (97–101)]. This may have left readers with the impression that CMVs are not controlled by the immune system.

This view, however, conflicts with the undisputable fact that acute CMV infections are rapidly and tightly controlled by the immune system, with CD8 T cells identified as being the main antiviral effector cells that terminate productive acute infection and surveil latent infection for preventing virus reactivation [(4,7,71,102), for a recent review see (103)]. Accordingly, CMVs do not harm the immunocompetent host but cause severe and often lethal, tissue-destructing organ infection in the immunocompromised host. This fundamental truth applies to humans as well as in the mouse model [reviewed in (2,3,6,9)].

CMVs are host-species specific, and different CMV species share homologous genes as well as genes with analogous function, but also possess ‘private genes’ that have evolved for adaptation to the respective host in eons of virus-host co-evolution. So, the evolutionary acquisition and maintenance of a gene involved in limiting direct viral antigen presentation in the MHC-I pathway to host CD8 T cells must be expected to have a benefit for both sides in the virus-host contract for co-existence. One hypothesis was that limiting direct antigen presentation to host CD8 T cells dampens the immune response to avoid virus clearance and to allow the virus to reach cellular niches for surviving in a state of latency. Our data now support a completely new view on the role of viral immune evasion.

It was our original aim to identify the canonical pathway of viral antigen presentation to CD8 T cells, that is, to decide between direct antigen presentation and cross-presentation. Previous studies in mouse models of CMV infection have shown that mice can mount a virus-specific CD8 T-cell response as a ‘plan B’ by either pathway when the respective other pathway is closed (51,54,55,104). Specifically, as also shown here, Unc93b1^3d/3d^ mice genetically deficient in cross-presentation can mount a CD8 T-cell response by direct presentation, and mice immunized with infected MHC-I-deficient cells can mount a CD8 T-cell response by cross-presentation (53). However, it remained open to question which pathway is used as ‘plan A’ when both pathways are accessible.

To tackle the problem, we used the novel approach of keeping the host immunogenetics constant and, instead, to genetically modify the virus in its immune evasion potential enhancing and reducing direct antigen presentation by infection with virus mutants mCMV-Δm152 and mCMV-m152.IE+E, respectively. If priming of naïve virus-specific CD8 T cells and subsequent clonal expansion are by direct antigen presentation, the magnitude of the CD8 T-cell response should have been high in case of infection with mCMV-Δm152, with which direct antigen presentation is not inhibited, and low in case of infection with mCMV-m152.IE+E, with which direct antigen presentation is strongly inhibited. The result was challenging: the response magnitude was the same for the two extremes of direct antigen presentation, seemingly proving that direct antigen presentation has no relevance at all for determining the magnitude of the CD8 T-cell response. What prevented us from drawing this rash conclusion was the curious result that intermediate immune evasion, and thus intermediate inhibition of direct antigen presentation, by infection with mCMV.WT led to the best response.

Notably, our current results for the response magnitude after infection with mCMV-Δm152 compared to mCMV.WT reproduce our previous findings, that is, reduced response after infection with a vRAP deletion virus mCMV-ΔvRAP despite better direct antigen presentation (69). We then explained this ‘immune evasion paradox’ of a low net response magnitude by a ‘negative feedback’ control of intranodal viral replication effected by the directly primed CD8 T cells, which limits the amount of antigen available for cross-presentation (69).

With this knowledge in mind, we herein analyzed intranodal viral replication after infection with all three viruses, that is, including now the new virus mCMV-m152.IE+E. The data again reproduced our previous findings for mCMV-ΔvRAP compared to mCMV.WT (69), but with the notable additional message that the highest intranodal replication after infection with mCMV-m152.IE+E, providing the highest amount of antigen for potentially feeding the cross-presentation pathway, did not generate a high net response magnitude. The intranodal replication of mCMV-m152.IE+E after infection is probably further enhanced by m152 interference with NKG2D-mediated NK cell activation (44,45) and suppression of type I interferon induction by STING (46). Thus, all functions of m152 support the supply with antigenic material for a potential cross-presentation. It is all the more surprising that such optimal conditions for cross-presentation did not enhance the CD8 T-cell response. Hence it follows that our own previous postulate of cross-presentation driving clonal expansion after initial priming by direct presentation (69) needs to be revised.

In conclusion, all available data for BALB/c mice now support the view that both, the initial event of priming naïve CD8 T cells as well as the subsequent clonal expansion, are driven by direct antigen presentation. The sketch in Fig. 7 summarizes the results for the example of BALB/c mice and illustrates the proposed mechanisms.

**Fig. 7:**
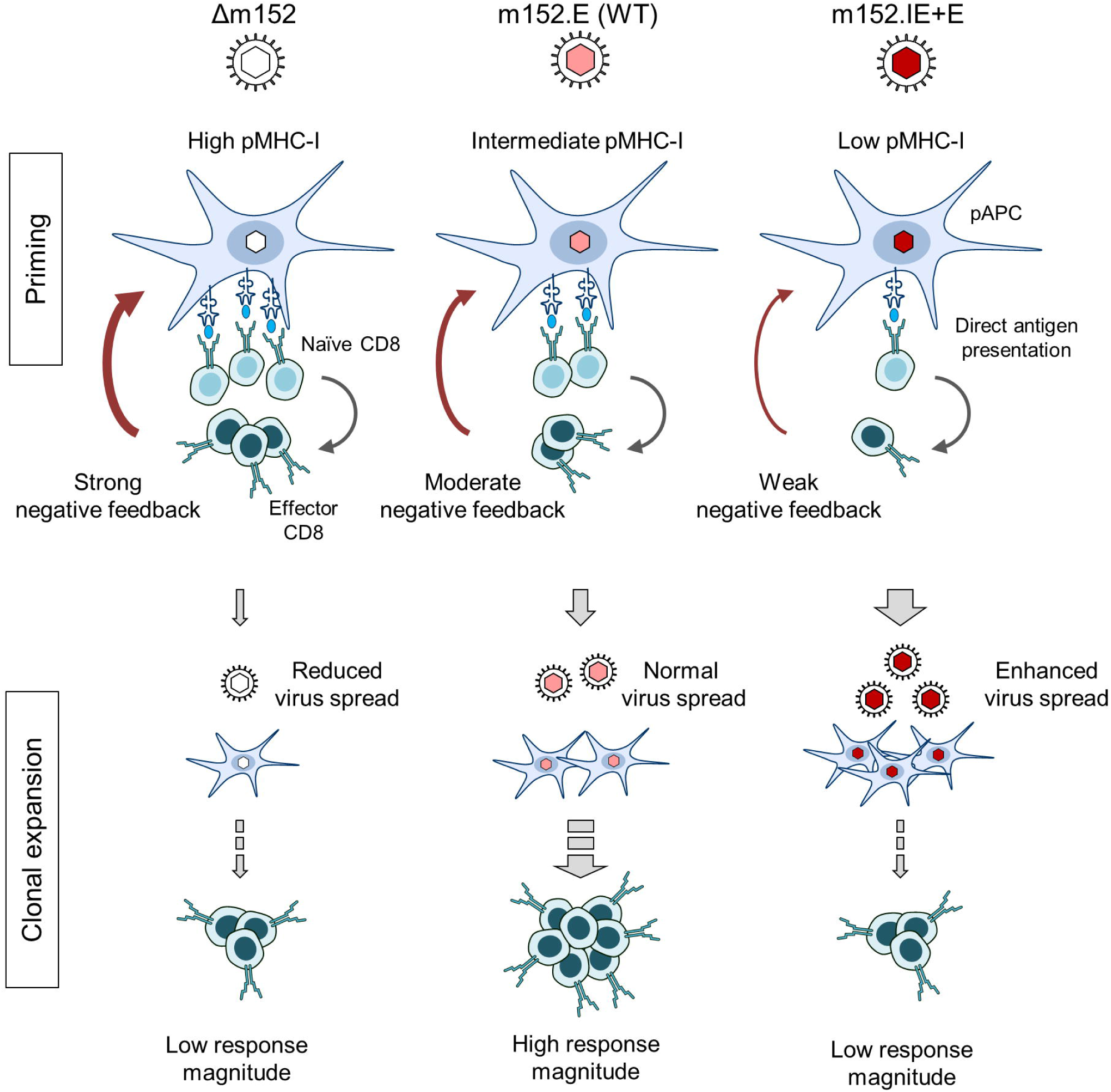
Graphical Abstract. Immunoevasin-dependent CD8 T-cell response magnitude regulated by negative feedback on direct antigen presenting cells by recently primed effector CD8 T cells. (pAPC), professional antigen-presenting cells; (pMHC-I), MHC-I molecules presenting antigenic peptides. Symbols on CD8 T cells represent viral epitope-specific T-cell receptors. Growing red-color intensity symbolizes increasingly enhanced expression of immunoevasin m152, and thus increasing strength of immune evasion.

Our data for mice with the C57BL/6 genetic background essentially support these principles. Like in BALB/c mice, the CD8 T-cell response to infection was not improved by an enhanced direct antigen presentation following the infection with mCMV-Δm152 compared to infection with mCMV-WT, with the only notable difference that it was not significantly reduced (recall Figs. 1A and 2A1). This finding is in accordance with an earlier observation made by Munks and colleagues (105), who concluded that the CD8 T-cell response to mCMV in C57BL/6 mice is not at all affected by immunoevasins. Our data, however, revealed that the net outcome is strongly influenced by activation of the Ly49H^+^ subset of NK cells via Ly49H-m157 ligation, a mechanism that accounts for genetic resistance against mCMV of mice with C57BL/6 genetic background, but is not operative in susceptible mouse strains, such as BALB/c [for a review, see (106)]. In accordance with direct antigen presentation, prevention of Ly49H^+^ NK-cell activity by either pan-NK cell depletion (Fig. 2B) or deletion of the viral ligand m157 (Fig. 3A) led to an enhanced CD8 T-cell response, despite reduced viral replication, in the RLN of mice infected with m152-deficient compared to m152-efficient virus. Nonetheless, a CD8 T-cell dependence of intranodal virus replication exists also in C57BL/6 mice (Figs.2B2, 2C and Figs. 3A2, 3B). We conclude that masked by activation of Ly49H^+^ NK cells, the negative feedback regulation by CD8 T cells in genetically mCMV-resistant C57BL/6 mice is weaker than in genetically mCMV-susceptible BALB/c mice, in which antiviral control is based primarily on CD8 T cells. This explains why negative feedback regulation after deletion of immunoevasin m152 only prevents enhancement of the CD8 T-cell response in C57BL/6 mice, whereas it reduces the CD8 T-cell response in BALB/c mice.

While we have now identified direct antigen presentation as the canonical pathway of antigen presentation for mounting a primary CD8 T-cell response in the ‘immunocompetent mouse’ model of CMV infection, in both a genetically-resistant and in a genetically-susceptible mouse strain, the findings have an even more important bearing for our understanding of viral interference with the MHC-I pathway of direct antigen presentation. We find it utmost intriguing that mCMV.WT, the virus naturally selected during virus-host co-evolution, induced the best CD8 T cell-response in genetically-susceptible BALB/c mice, whereas both prevention and enhancement of m152-mediated immune evasion diminished the response. It appears that optimal calibration of the strength of viral interference with the MHC-I pathway of antigen presentation serves to still allow sufficient priming of naïve CD8 T cells to initiate a response but also to moderate the ‘negative feedback’ that otherwise would inhibit clonal expansion of the primed CD8 T cells. So, unexpectedly, viral interference with direct antigen presentation is beneficial for mounting a protective CD8 T-cell response. This sheds a completely new light on the physiological role of viral immune evasion.

## Acknowledgements

The authors thank Angélique Renzaho and Kirsten Freitag for expert technical assistance and Daria Lohschelders for her help during manuscript preparation.

## Biosafety Statement

The work was done according to German federal law GenTG and BioStoffV. The generation of recombinant mCMV was approved by the ‘Struktur-und Genehmigungsdirektion Süd’ (SDG, Neustadt, Germany), permission numbers 24.1-886.3.

## Data Availability Statement

The original contributions presented in the study are included in the article/Supplementary Material. Further inquiries can be directed to the corresponding author.

## Ethics Statement

The animal study was reviewed and approved by the ethics committee of the ‘Landesuntersuchungsamt Rheinland-Pfalz’ according to German federal law §8 Abs. 1 TierSchG (animal protection law), permission numbers 177-07/G09-1-004.

## Author Contributions

MJR initiated the project. MJR and NAL are responsible for conception and design of the study, analysis, and interpretation of the data. RH is responsible for parts of data analysis and interpretation. MMB provided essential materials and designed parts of the study. JKB, SB, AF, and NAL conducted the work and analyzed the data. NAL and MJR wrote the manuscript. All authors contributed to manuscript revision and read and approved the submitted version.

## Figure Legends

**Suppl. Fig. 1:** Magnitude and specificity repertoire of the acute CD8 T-cell response response in BALB/c mice. Frequencies of CD8 T cells responding to infection with either mCMV.WT (WT, two upper panels) or mCMV-m152.IE+E (m152.IE+E, two lower panels) were determined by intracellular IFNγ-staining, using as stimulator cells either an mCMV genome-wide ORF library of transfectants (75) (left panels) or P815 cells exogenously-loaded with the indicated synthetic antigenic peptides at the saturating concentration of 10^−7^ M (right panels). Responder cells were splenocytes isolated on day 7 after intraplantar infection with 1×10^5^ PFU each of either of the two viruses. Note that the comparison of ORF library data for mCMV.WT and mCMV-ΔvRAP, which is equivalent to mCMV-Δm152, has been published previously (107), and supports our conclusions.

**Suppl. Fig. 2:** Viral replicative fitness in host organs. Immunocompromised BALB/c mice (6.5 Gy of total body γ-irradiation) were infected with 1×10^5^ PFU of mCMV.WT (WT), mCMV-m152.IE+E (m152IE+E) or mCMV-Δm152 (Δm152). Viral replicative fitness was assessed by the viral doubling times (vDT), measured by M55/gB-specific qPCR in total DNA extracted from the organs indicated. Symbols represent individual mice. vDT values and their 95% confidence intervals were calculated from log-linear regression lines with ordinate intercept, including all data collected over the entire time course. Dashed curves border the respective 95% confidence areas.

