## Supplementary figures and images for "Direct Antigen Presentation is the Canonical Pathway of Cytomegalovirus CD8 T-cell Priming Regulated by Balanced Immune Evasion Mounting a Strong Antiviral Response"

### Suppl. Figure 1

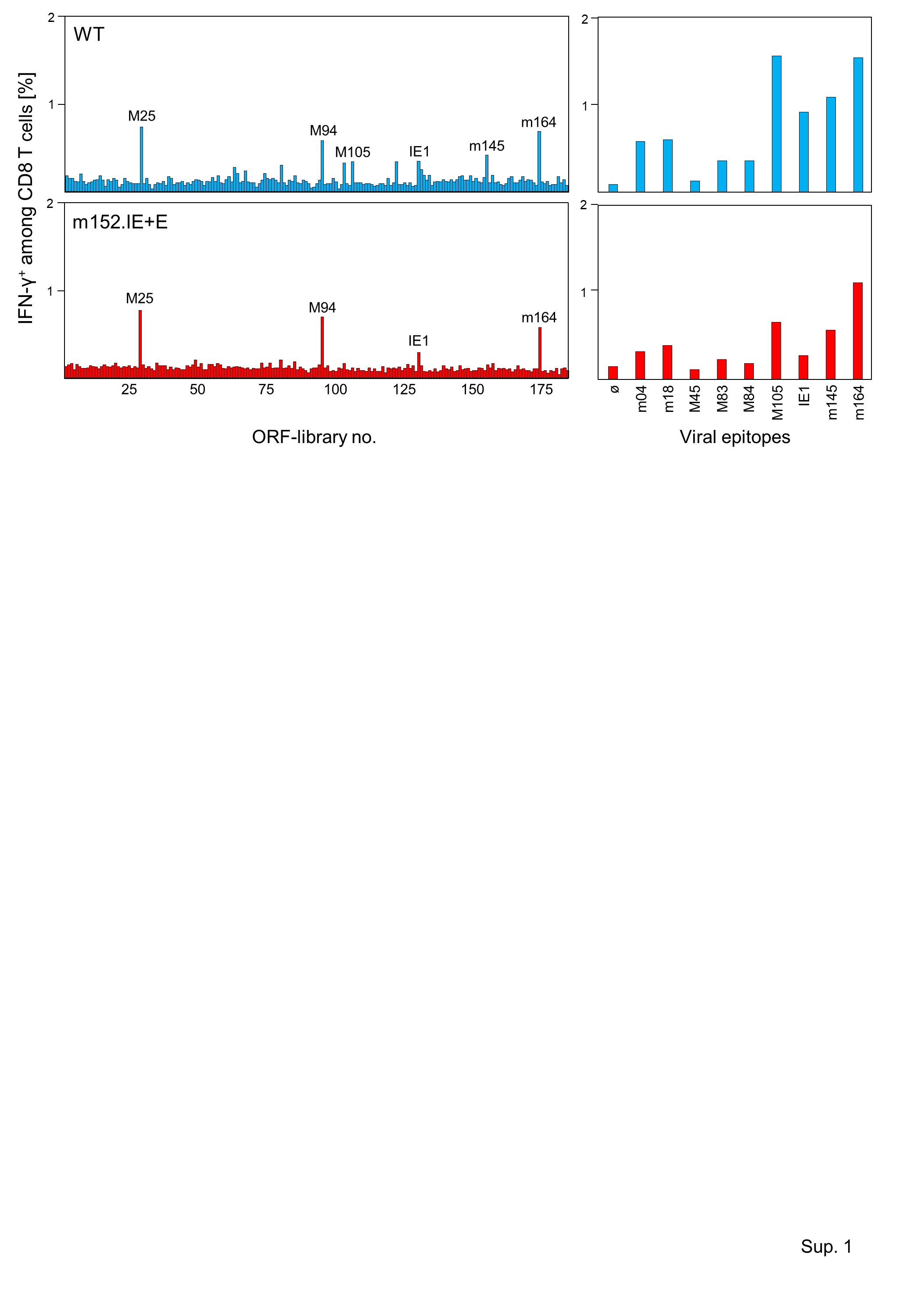

### Suppl. Figure 2

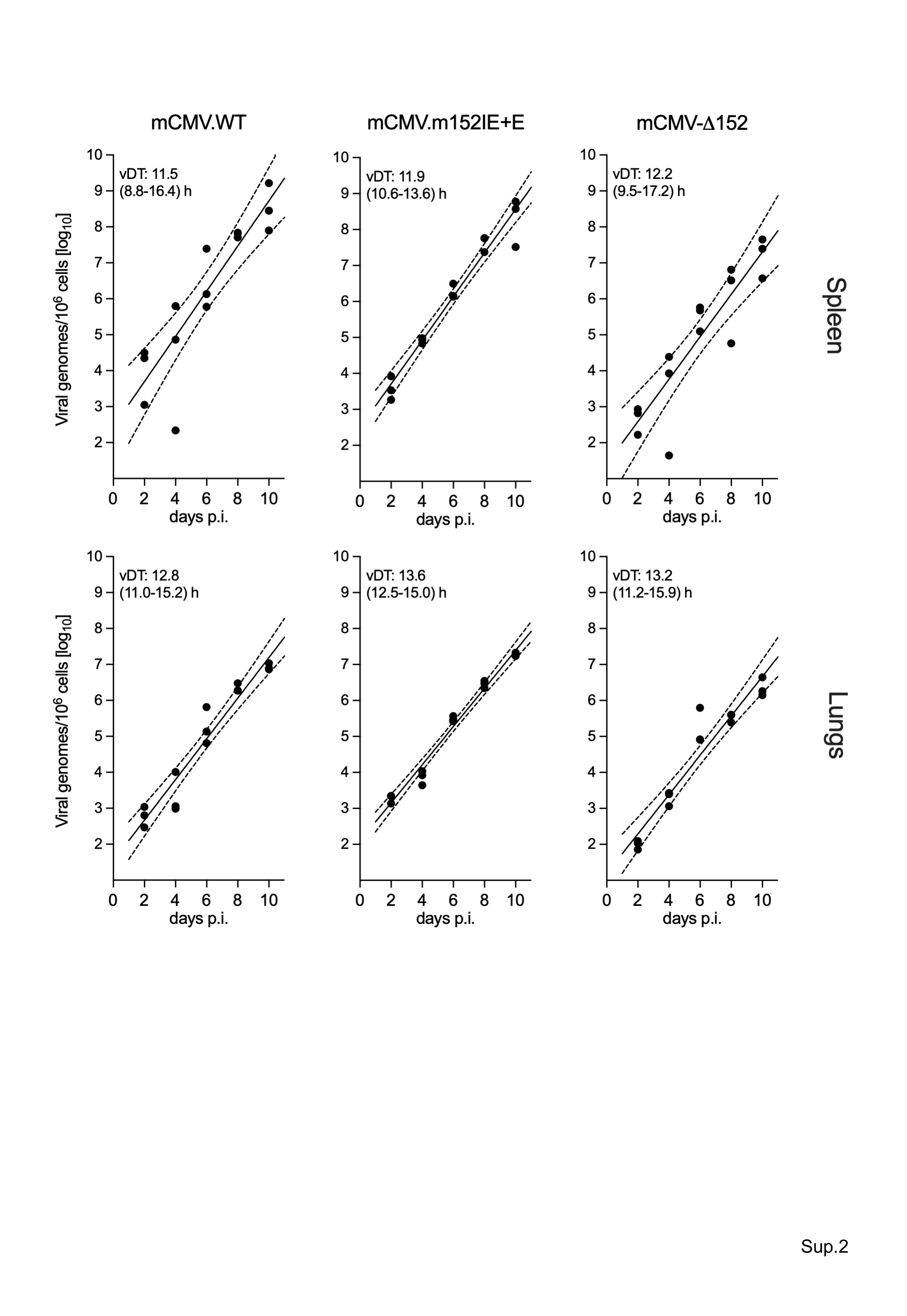
